# Large-scale insights into the biosynthetic potential of the *Bacillus cereus* group

**DOI:** 10.1101/2025.10.16.682773

**Authors:** Josefin Blom, Joseph Wambui, Hadrien Gourlé, Martin Larralde, Vignesh Ramnath, Johan Henriksson, Laura M. Carroll

**Affiliations:** Department of Clinical Microbiology, SciLifeLab, Umeå University, Umeå, Sweden; Laboratory for Molecular Infection Medicine Sweden (MIMS), Umeå University, Umeå, Sweden; Umeå Centre for Microbial Research (UCMR), Umeå University, Umeå, Sweden; Institute for Food Safety and Hygiene, University of Zürich, Zürich, Switzerland; Structural and Computational Biology Unit, European Molecular Biology Laboratory, Heidelberg, Germany; Leiden University Center for Infectious Diseases (LUCID), Leiden University Medical Center, Leiden, Netherlands; Department of Molecular Biology, SciLifeLab, Umeå University, Umeå, Sweden; Integrated Science Lab (IceLab), Umeå University, Umeå, Sweden

## Abstract

Bacterial secondary metabolites (SMs) are a critical source of natural product-derived drugs. However, SM discovery efforts have focused overwhelmingly on *Actinomycetes*, potentially overlooking other key producers. Here, we explore the biosynthetic potential of the *Bacillus cereus* group, an underexplored complex of SM producers. Using a combined rule- and machine learning-based approach, we mine an unprecedented number of *B. cereus* group genomes (*n* = 9,744) for SM-producing biosynthetic gene clusters (BGCs; *n* = 200,196). Notably, 158,678 *B. cereus* group BGCs (78.2%) did not cluster with previously described BGCs, suggesting new chemical scaffolds to be explored. *B. pseudomycoides* was particularly prolific in terms of its SM production potential (30.8 BGC families/genome, Kruskal-Wallis *p* < 0.0001), and we identify a previously uncharacterized, *B. pseudomycoides*-unique peptide. Overall, our study represents the largest survey of *B. cereus* group biosynthetic potential to date and posits the complex as an under-queried SM resource.

## INTRODUCTION

Secondary metabolites (SMs; also known as specialized metabolites or natural products) are non-essential molecules employed by bacteria for selective advantages in their ecological niche (Tyc *et al*. 2017; Mullowney *et al*. 2023). As crucial components of bacterial inter- and intraspecies interplay in complex environments, SMs allow their producers to respond to stressors and stimuli, scavenge nutrients and raise their stress tolerance (Tyc *et al*. 2017; Milshteyn, Colosimo and Brady 2018; Perry, Meirelles and Newman 2022; Mullowney *et al*. 2023). Beyond their ecological importance, bacterial SMs are a valuable resource of chemical scaffold diversity, which has been used to develop a wide range of products with important industrial applications (Medema and Fischbach 2015; Atanasov *et al*. 2021; Mullowney *et al*. 2023). Bacterial SMs have been especially important within the realm of drug discovery, with approximately 50% of clinically used antibiotics derived from the *Streptomyces* genus (Tyc *et al*. 2017; Parra *et al*. 2023).

With the bulk of previously explored SMs originating from a few bacterial species, it has been suggested that only a limited fraction of microbial biosynthetic capacity has been unveiled to date (Doroghazi *et al*. 2014; Medema and Fischbach 2015; Hemmerling and Piel 2022; Parra *et al*. 2023). Past research has demonstrated that microbes present in complex, free-living environments with high degrees of competition frequently produce high numbers of SMs (Machado *et al*. 2021), and therefore ought to be prioritized in investigative efforts. The *Bacillus cereus* group (*B. cereus sensu lato*; *B. cereus s*.*l*.) is one such group of microbes, as members of this Gram-positive, spore-forming species complex are nearly ubiquitous in nature, particularly within soil and marine environments (Ehling-Schulz *et al*. 2006; Stenfors Arnesen, Fagerlund and Granum 2008; Mondol, Shin and Islam 2013). Several SMs produced by *B. cereus* group members (e.g. zwittermicin A, cerexin A, pseudomycoidicin) have already been shown to have potential as drug candidates, given their antimicrobial activity against Gram-positive bacteria (Shoji and Hinoo 1975; Shoji, Kato and Sakazaki 1976; Kevany, Rasko and Thomas 2009; Basi-Chipalu *et al*. 2015). Conversely, other SMs produced by *B. cereus* group members are known to play negative roles in human health.

Cereulide, for example, is a cyclic dodecadepsipeptide which causes emetic foodborne illness in humans (Stenfors Arnesen, Fagerlund and Granum 2008; Ehling-Schulz, Frenzel and Gohar 2015; Morandini *et al*. 2024). Recently, bacillamide D, a non-ribosomally produced peptide (NRP) produced by a *B. cereus* group strain isolated from the mouse intestine, was shown to be highly cytotoxic to multiple human cell lines (Lagkouvardos *et al*. 2016; Hohmann *et al*. 2024). Consequently, the (i) environmental ubiquity and (ii) known ability to produce SMs of medical/industrial relevance make the *B. cereus* group a promising candidate reservoir for novel SMs.

Previous attempts to assess the biosynthetic potential of the *B. cereus* group (Zhao and Kuipers 2016; Grubbs *et al*. 2017; Xia *et al*. 2022; Vater *et al*. 2023; Yin *et al*. 2023) have made use of the fact that the biosynthetic pathways responsible for SM production are typically encoded by physically clustered groups of genes called biosynthetic gene clusters (BGCs) (Medema and Fischbach 2015). In each of these earlier studies, *B. cereus* group genomes were mined for BGCs; however, they relied on a limited number of *B. cereus* group genomes (i.e., no more than 3,381 genomes at most) and/or did not include all taxa within the group (Zhao and Kuipers 2016; Grubbs *et al*. 2017; Xia *et al*. 2022; Vater *et al*. 2023; Yin *et al*. 2023). Further, prior research studying the *B. cereus* group’s biosynthetic potential relied solely on rule-based approaches (e.g., antiSMASH, BAGEL), which use hard-coded “rules” for BGC detection (Zhao and Kuipers 2016; Grubbs *et al*. 2017; Van Heel *et al*. 2018; Xia *et al*. 2022; Blin *et al*. 2023; Mullowney *et al*. 2023; Vater *et al*. 2023; Yin *et al*. 2023). However, recent bioinformatic advances have resulted in machine learning (ML) methods, which can detect BGCs of novel architecture (Cimermancic *et al*. 2014; Hannigan *et al*. 2019; Carroll *et al*. 2021; Mullowney *et al*. 2023). Whether ML-based methods enable the discovery of novel BGCs compared to rule-based methods alone remains unknown within the context of the *B. cereus* group.

Here, we mine an unprecedented number of *B. cereus* group genomes for BGCs, with the goal of providing large-scale insights into the biosynthetic potential of the species complex. Using a combined rule- and ML-based approach, we detect >200,000 BGCs in >9,700 *B. cereus* group genomes, the majority of which are putative novel BGCs. Using our compendium of BGCs, we identify taxa within the *B. cereus* group which harbor significantly more BGCs than other members of the species complex (i.e., SM “superproducers”). Finally, to make our predicted BGCs accessible to the public, we present BTyperBGC (https://github.com/c20josbl/btyperbgc_shiny), our interactive *B. cereus* group BGC atlas, which links SM-associated data to manually curated, standardized metadata to enable SM discovery. Overall, our study represents the largest survey of *B. cereus* group biosynthetic potential to date and demonstrates that the *B. cereus* group is an under-queried reservoir of putative novel SMs.

## RESULTS

### BTyperBGC enables large-scale BGC exploration within the *B. cereus* group

To create BTyperBGC, our interactive *B. cereus* group BGC atlas, we mined all high-quality *B. cereus group* genomes from BTyperDB (*n* = 9,744; Ramnath *et al*. 2023) for BGCs, using two approaches: (i) antiSMASH, a popular and precise rule-based method (Blin et al. 2021); (ii) GECCO, a ML-based method which prioritizes BGC novelty (Figure 1; Carroll et al. 2021). In total, our combined approach identified 200,196 BGCs, of which 104,847 and 95,349 were detected using antiSMASH and GECCO, respectively (Supplementary Table S1, Supplementary Figure S1). The resulting *B. cereus* group BGCs were combined with the Minimum Information about a Biosynthetic Gene Cluster (MIBiG) database (v4.0; *n* = 2,636 BGCs; Zdouc et al. 2025), a curated database of previously identified and characterized BGCs, to enable comparisons between the newly mined *B. cereus* group BGCs and those described in literature. To reduce redundancy, remove duplicate BGCs and enable functional prediction via comparison to the MIBiG BGCs, all BGCs (*n* = 202,832; referred to hereafter as the “full BGC set”) were clustered into gene cluster families (GCFs, *n* = 2,629 GCFs; Supplementary Table S1). For two-dimensional visualization of the dataset, representative BGCs (one BGC from each GCF) were annotated using the Pfam database, and the resulting protein domain composition matrix was used to create a UMAP (see sections “Protein domain annotation” and “UMAP construction” below for details), forming the basis of BTyperBGC. BTyperBGC provides an overview of the biosynthetic space of the *B. cereus* group and can be annotated and filtered based on genomic and metadata features to facilitate selection of experimental leads for further testing. Examples of filters include several standardized taxonomic nomenclatures commonly used in *B. cereus* group research, e.g. *panC* group assignments (Guinebretière *et al*. 2008, 2010; Carroll, Cheng and Kovac 2020), the Genome Taxonomy Database (GTDB) taxonomy (Parks *et al*. 2022), and the 2020 *B. cereus* group Genomospecies-Subspecies-Biovar (GSB) taxonomy implemented in BTyper3 (Carroll, Cheng and Kovac 2020; Carroll, Wiedmann and Kovac 2020). For the remainder of this study, the BTyper3 *panC* nomenclature will be used unless otherwise stated, as it is likely the most interpretable to readers.

**Figure 1.**
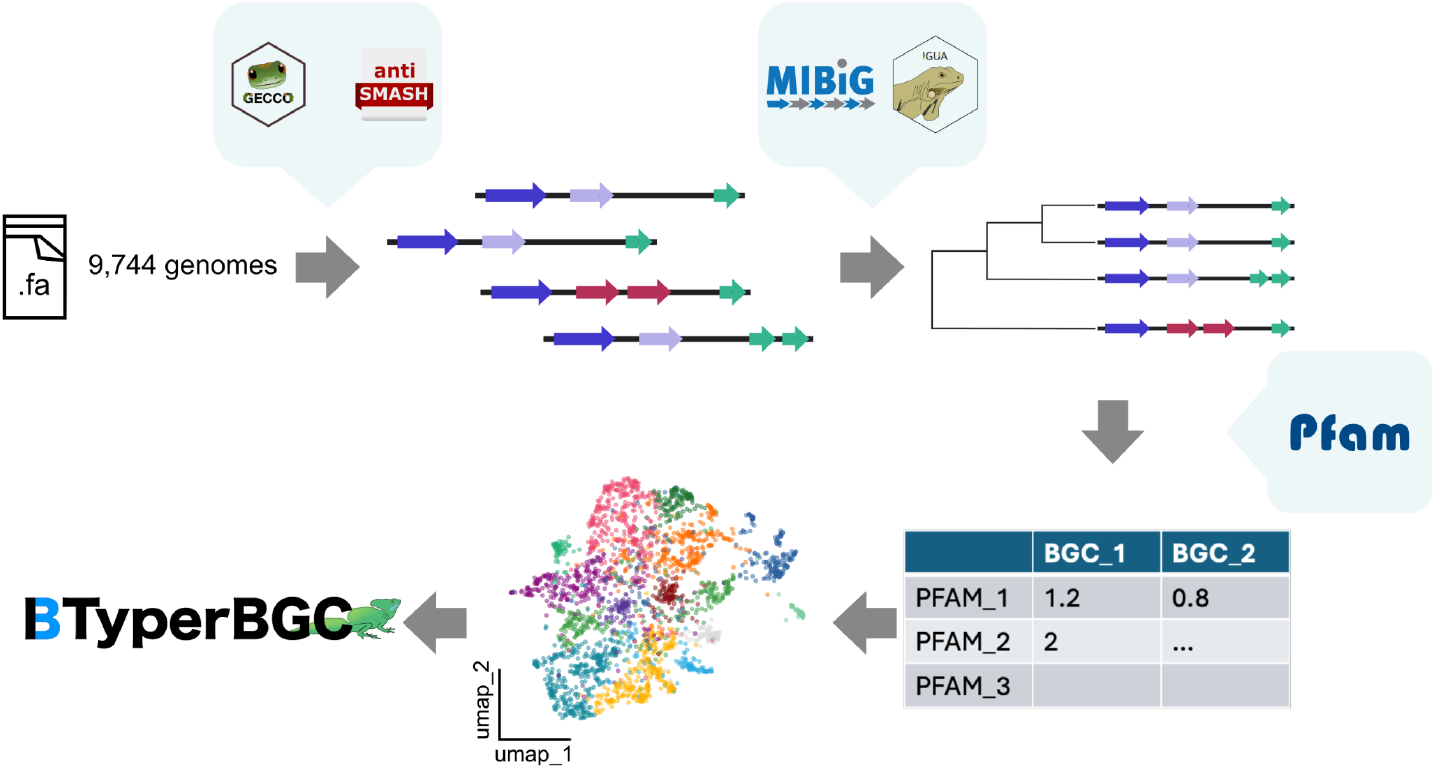
An overview of the construction of BTyperBGC. Briefly, two BGC detection tools (antiSMASH and GECCO) were used to identify biosynthetic gene clusters (BGCs) in 9,744 *B. cereus* group genomes. The resulting BGCs (*n* = 200,196) were clustered together with previously known BGCs from the MIBiG database to form gene cluster families (GCFs; *n* = 2,629). GCF representative BGCs (one per GCF) were annotated using Pfam, and the resulting protein domain composition matrix was used to create a UMAP (i.e., the two-dimensional visualization underlying BTyperBGC).

### A large proportion of *B. cereus* group biosynthetic gene cluster families represent putative novel BGCs

In comparison to well-known SM producers such as *Streptomyces*, genomic mining efforts aimed at the *B. cereus* group have been sparse. We therefore hypothesized that a vast fraction of BGCs encoded by the *B. cereus* group is yet to be discovered and/or characterized. Using the BGCs from BTyperBGC, the majority of GCFs contained no *B. cereus* group BGCs (i.e., MIBiG BGCs alone; 62.9%, *n* = 1,653 GCFs), and were thus excluded from further analysis. Of the remaining GCFs, only 41 contained a mixture of MIBiG BGCs (*n* = 62 BGCs) and BGCs derived from *B. cereus* group genomes (*n* = 41,580 BGCs; Figure 2). After disregarding MIBiG BGC BGC0002899 (currently pending in MIBiG), further examination of the MIBiG-containing GCFs revealed that the largest four encompassed the MIBiG entries petrobactin, a *Bacillus anthracis* siderophore required for virulence in mice (Cendrowski, MacArthur and Hanna 2004), and an array of BGCs known to have antimicrobial activity (i.e., thurincin H, thuricin, zwittermicin A and thusin) (Rea *et al*. 2010; Wang *et al*. 2014; Xin *et al*. 2016; Ortiz-Rodríguez *et al*. 2023). Notably, the relevant MIBiG entries originated among *B. cereus* group members, and therefore perhaps a natural outcome that related compounds were seemingly detected among our full BGC dataset (Figure 2C).

**Figure 2.**
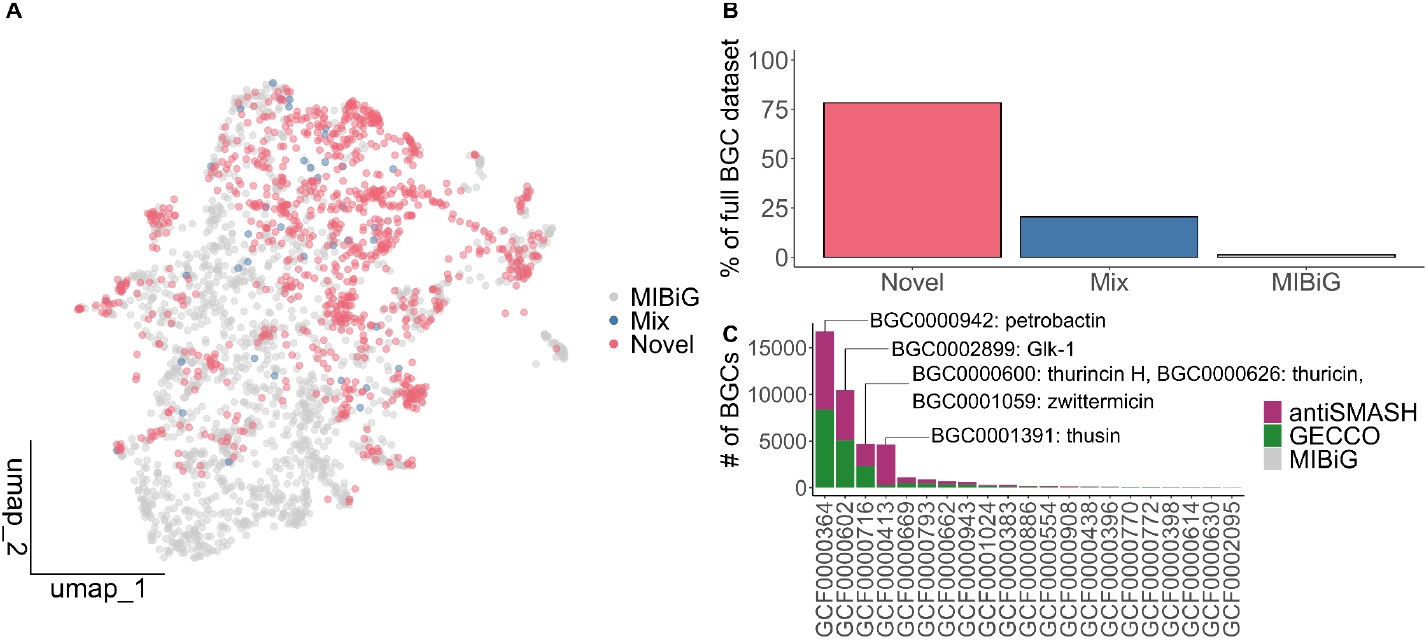
(A) UMAP of gene cluster family (GCF) representative BGCs (points) colored by whether a GCF contained (i) solely MIBiG BGCs (MIBiG), (ii) BGCs mined by antiSMASH and/or GECCO (Novel), or (iii) BGCs from MIBiG and GECCO/antiSMASH (Mix). (B) Bar plot showing the percentage of BGCs within the full BGC dataset (y-axis; *n* = 202,832 total BGCs) belonging to the Novel, MIBiG and Mix categories (x-axis), as described for (A). (C) Stacked bar plot demonstrating the origin of BGCs present in the Mix GCFs. For the four largest GCFs, the MIBiG BGCs (v4.0) have been highlighted. Mix GCFs with less than 20 BGCs have been removed.

Intriguingly, the bulk of *B. cereus* group-derived BGCs were contained within GCFs devoid of MIBiG BGCs (*n* = 158,678 BGCs, 78.2% of the full BGC dataset; Figure 2B). A substantial proportion of these 935 GCFs (46.2%, *n* = 432 GCFs) contained exclusively GECCO-derived BGCs, compared to GCFs which were antiSMASH-exclusive or included a mixture of GECCO- and antiSMASH-derived BGCs (12.1% and 41.7%, respectively; Supplementary Figure S3). Thus, the ML-based BGC mining method captured GCFs not detected using the rule-based alternative, thereby providing support for our combined approach.

### Biosynthetic gene cluster families are largely species- or strain-specific within the *B. cereus* group

Members of the *B. cereus* group share extensive parts of their genetic make-up (Rasko *et al*. 2005; Ehling-Schulz, Lereclus and Koehler 2019). Therefore, we investigated whether the potential to synthesize SMs was shared or distinct among genomes and species (here, *panC* groups). In total, 22.6% of GCFs were unique to a single genome (*n* = 221 GCFs, of which 168 GCFs were singleton GCFs), and the majority of GCFs (52.6%, *n* = 513 GCFs) were confined to <5 genomes (Supplementary Figure S2).

Moreover, no GCFs were common to all *B. cereus* group genomes analyzed here (Supplementary Table S1, Supplementary Figure S2), and only 10 GCFs (1.02% of 976 GCFs) encompassed BGCs from over half of the included genomes (>4,872 genomes). However, GCFs GCF0000534, GCF0001105 and GCF0000586 contained BGCs from >9,600 genomes (99.9%, 99.9% and 99.0%, respectively), and were therefore highly prevalent in our dataset. For these three GCFs, the majority biosynthetic class was “Others”. Similarly, while three GCFs spanned all *panC* groups (GCF0000369, GCF0000605 and GCF0001105), nearly half of all GCFs were confined to a single *panC* group (462 of 976 GCFs, 47.3%; Supplementary Figure S2). Neither the GCFs that spanned the maximum number of genomes, nor those encompassing all *panC* groups contained any BGCs from MIBiG (Supplementary Table S1). To conclude, while some *B. cereus* group GCFs are widespread throughout the group, most GCFs are comparatively strain-specific (i.e., confined to a very few genomes) and/or species-specific (i.e., confined to a single *panC* group; Supplementary Figure S2).

### *B. pseudomycoides* presents as a potentially prolific secondary metabolite producer among the *B. cereus* group members

Given the relative strain-specificity of *B. cereus* group GCFs, we asked whether certain *B. cereus* group species (here, *panC* groups) encoded unusually large numbers of BGCs (Figure 3). As two BGC detection methods were used, the full BGC dataset likely contained duplicate BGCs. Thus, to avoid counting redundant BGCs, we used the number of GCFs per genome as a proxy for biosynthetic potential, premised on the assumption that GCFs represent unique chemical scaffolds (Navarro-Muños *et al*. 2020). Furthermore, we excluded *panC* groups with fewer than five genomes, hoping to avoid biased means due to few data points (i.e., genomes). Among the remaining *panC* groups, *B. pseudomycoides* (*panC* Group I) had the highest mean GCFs per genome (30.8 GCFs per genome, *n* = 153 genomes; Kruskal-Wallis test: *H*(8) = 5,796, *p* < 0.0001; Dunn’s Test with Holm correction: *Z* = 7.3709944-27.8542690, *p adj*. < 0.0001).

**Figure 3.**
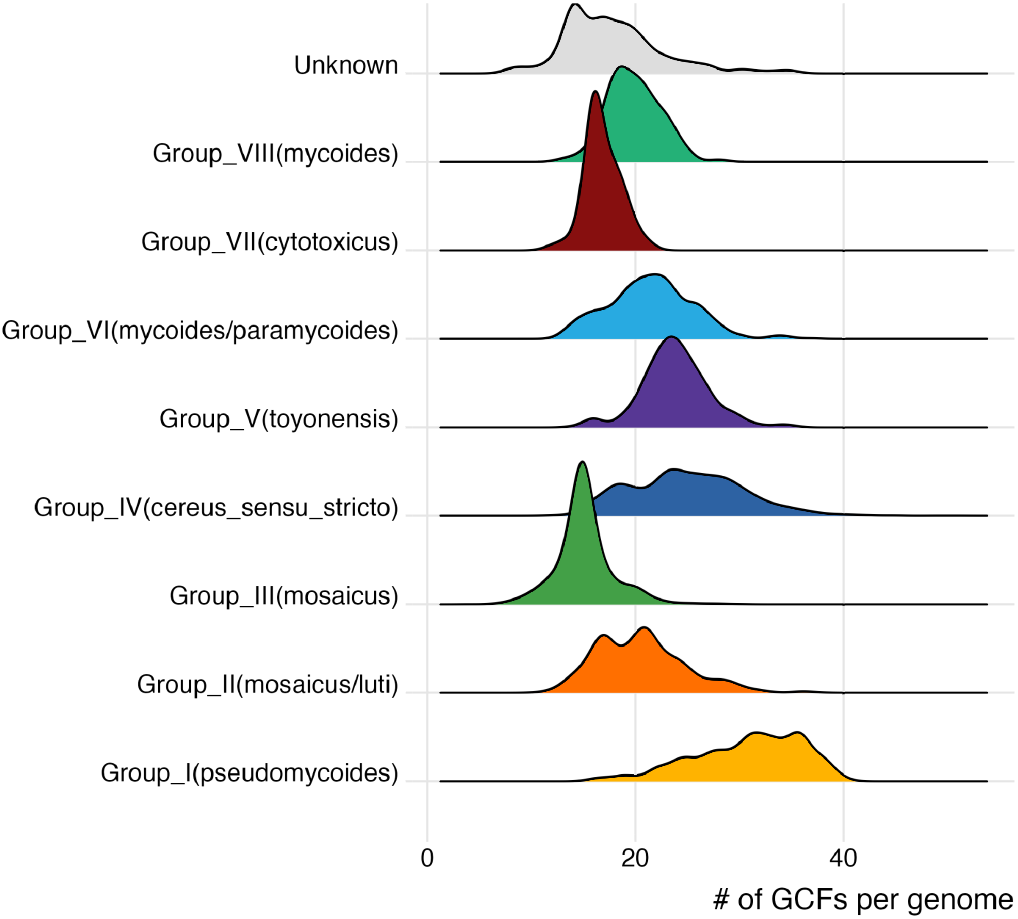
Ridge plot of gene cluster families (GCFs) per genome (x-axis) per *panC* group (y-axis).

As (i) *panC* Group I (i.e., *B. pseudomycoides*) presented with an unusually high number of GCFs per genome, and (ii) *B. cereus* group BGCs were largely novel, we decided to investigate lineage-associated patterns of GCFs unique to *panC* Group I (referred to hereafter as “*B. pseudomycoides*-unique GCFs”, *n* = 31 GCFs; Figure 4). Notably, the largest of the *B. pseudomycoides-*unique GCFs, GCF0001397 (*n* = 100 BGCs), seemed comparatively clade-specific; using *in silico* seven-gene multi-locus sequence typing (MLST), this GCF included genomes of the following sequence types (STs): ST1346 (*n* = 11), ST2130 (*n* = 3), ST87 (*n* = 3), ST3093 (*n* = 1) and of unknown ST (*n* = 9;

**Figure 4.**
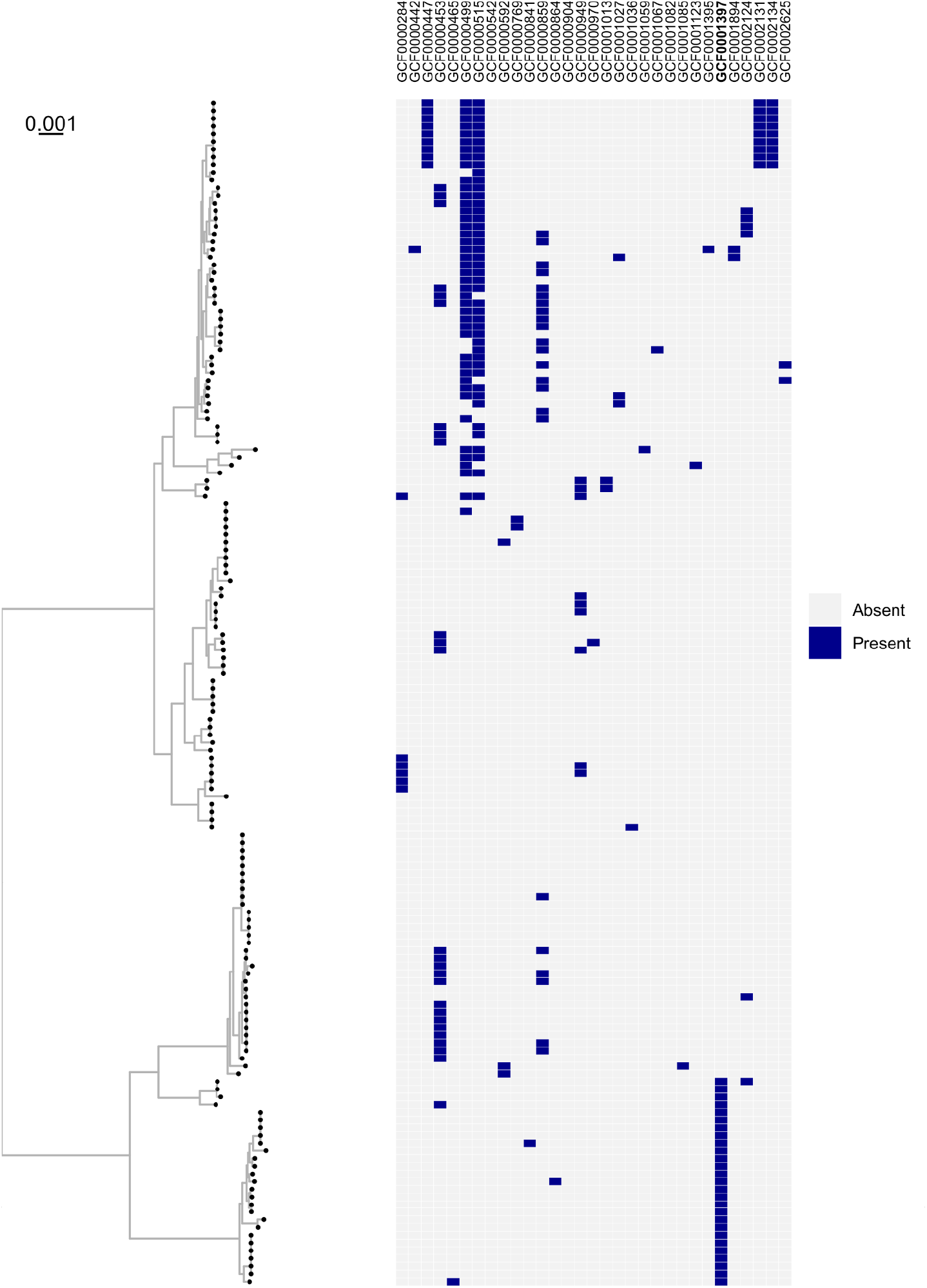
A maximum-likelihood phylogeny of genomes belonging to Genome Taxonomy Database (GTDB) species *B. pseudomycoides* (left) with annotation showing absence/presence of BGCs in *B. pseudomycoides*-unique gene cluster families (GCFs; right). The phylogeny is rooted via an outgroup (the *B. bingmayongensis* type strain genome, BTyperDB ID BTDB_2022-0000619.2, omitted for readability), with branch lengths reported in substitutions per site. GCF0001397 has been highlighted in bold font.

Supplementary Table S2). While other *B. pseudomycoides-*unique GCFs exhibited some level of clade-specificity, chief amongst them GCF0000499 and GCF0000515 (*n* = 43 and 44 BGCs, respectively; Figure 4), we chose to focus on GCF0001397, due to the relatively large numbers of BGCs it encompassed.

Most of the BGCs present in GCF0001397 (*n* = 100 BGCs) were predicted to be NRPs (96%, 96 BGCs), with a minority designated as ribosomal BGCs (4%, 4 BGCs; Supplementary Table S2). Additionally, while the BGCs varied in length and genomic content, the majority included an ABC transporter, likely for export of the final metabolite, and NRP synthetases (Figure 5, Supplementary Figure S4). The latter most closely matched NRP synthetases of the known BGCs sevadicin, mycosubtilin and bacillibactin (Figure 5). However, direct comparisons between the NRP synthetases of a selected GCF0001397 representative and the MIBiG BGCs for sevadicin, mycosubtilin and bacillibactin demonstrated that this similarity was rather limited (% amino acid identity = 19.1-37.8 for query coverages >90%, Figure 5), suggesting that the BGCs may produce a novel peptide uniquely present among a clade of *B. pseudomycoides* within the *B. cereus* group.

**Figure 5.**
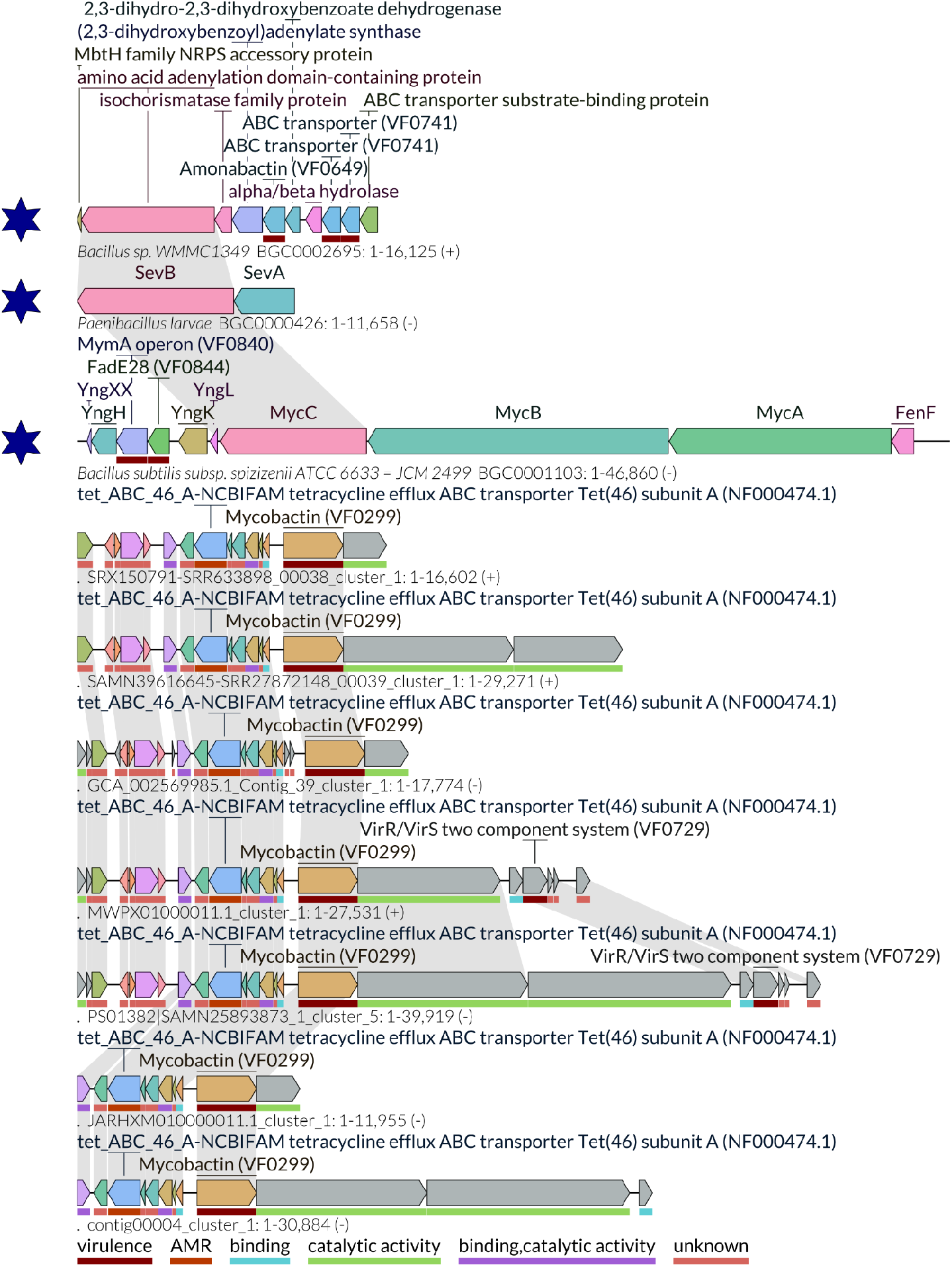
LoVis4u plot of a sub-sample of GCF0001397, together with BGC0000426 (sevadicin), BGC0001103 (mycosubtilin) and BGC0002695 (bacillibactin) from MIBiG (v4.0; dark blue stars). An equivalent plot including all GCF0001397 BGCs mined by GECCO can be found in the supplementary material (Supplementary Figure S4).

## DISCUSSION

Using prokaryotic SM chemical scaffold diversity to boost the number of leads in drug discovery pipelines has been presented as an alternative strategy to, e.g., *in silico* drug design (Atanasov *et al*. 2021; Hemmerling and Piel 2022). With the historic success of natural product-based drug design, there is an incentive to expand genomic mining efforts to under-utilized species (Scherlach and Hertweck 2021; Parra *et al*. 2023), such as the *B. cereus* group, to discover new classes of drugs. Here, we mined 9,744 *B. cereus* group genomes for BGCs, thereby tripling the number of genomes queried in previous studies of the *B. cereus* group’s biosynthetic potential (Zhao and Kuipers 2016; Grubbs *et al*. 2017; Xia *et al*. 2022; Vater *et al*. 2023; Yin *et al*. 2023). Further, our combined rule- and ML-based BGC mining strategy allowed us to explore the BGCs of this species complex beyond the confines of classic approaches, and in the process generate a comprehensive dataset of >200,000 BGCs. The expansiveness of the dataset supports the notion that *B. cereus* group members are comparatively prolific producers of SMs, albeit not to the extent of *Streptomyces* (Gavriilidou et al. 2022; Mohite et al. 2025).

Our research also puts forward *B. pseudomycoides* as a prominent SM “superproducer” within the *B. cereus* group. A prior study has hinted that *B. pseudomycoides* might possess a high number of BGCs, but only included a singular *B. pseudomycoides* strain (Grubbs *et al*. 2017). Additional investigation into *B. pseudomycoides*-derived BGCs has been scarce, with the one notable example being pseudomycoidicin, a class I lanthibiotic with lytic activity against Gram-positive bacteria (Basi-Chipalu *et al*. 2015). Here, we explored a seemingly clade-specific, *B. pseudomycoides-*unique GCF, which likely encodes NRPs with little to no similarity to previously described BGCs. These BGCs represent emergent natural product leads, which could be explored experimentally in the future.

Most importantly, the small intersection observed between the *B. cereus* group BGCs and those present in MIBiG indicates novel chemical scaffolds with unknown biological activity, which could be harnessed for the development of e.g., medically important therapeutics, entomopathogenic biopesticides and antifungals. To facilitate further development, testing and application of *B. cereus* group-derived metabolites, we created an online resource for the largest collection of *B. cereus* group-derived BGCs to date (BTyperBGC), through which the full BGC dataset can be explored in conjunction with genomic metadata, e.g. standardized species assignments. However, while *in silico* BGC mining is rapid and scalable, these methods generate vast quantities of predicted BGCs. Translating the predictions to beneficial applications requires high-throughput experimental testing, which despite recent developments (Meena *et al*. 2024; Yang *et al*. 2025), remains an unsurmounted challenge. Nevertheless, our research provides an important first step and initial insight into the possibilities of harnessing the *B. cereus* group species complex for natural product-based drug discovery on a large scale.

## METHODS

### BGC detection

*B. cereus* group genomes (*n* = 9,744) and associated metadata were acquired from BTyperDB (Ramnath *et al*. 2023; Supplementary Table S1; access date 31st of March, 2025). BGCs were detected in each genome using two methods: (i) antiSMASH (v6.1.1; Blin et al. 2021) and (ii) GECCO (v0.9.8; Carroll et al. 2021). For (i), antiSMASH was used to detect BGCs with the prodigal-m option (--genefinding_tool prodigal-m) enabled. Defaults were used for all remaining settings. For (ii), GECCO was run with the probability threshold lowered to 0.3 (to prioritize novelty detection; -m 0.3), the mask option (--mask) enabled, and defaults used for remaining settings. BGCs detected by antiSMASH were assigned the prefix of “AS.” (Supplementary Table S1), to allow for differentiation.

### BGC clustering

To compare *B. cereus* group BGCs detected *in silico* (see section “BGC detection” above) to previously described/characterized BGCs, all BGCs present in the MIBiG database (*n* = 2,636; v4.0; Zdouc et al. 2025) were downloaded. BGCs detected via (i) antiSMASH (*n* = 104,847) and (ii) GECCO (*n* = 95,349) were aggregated with (iii) BGCs from MIBiG to generate the full BGC dataset of 202,832 BGCs (Supplementary Table S1). The full BGC dataset was supplied as input to IGUA (v0.1.0; Larralde et al. 2025), which was used to cluster BGCs into GCFs using default settings. IGUA produced a total of 2,629 GCFs and a set of 2,629 GCF representative BGCs (i.e., one representative BGC per GCF, referred to hereafter as the “GCF representative set”; Supplementary Table S1).

### Protein domain annotation

BGCs in the GCF representative set, as selected by IGUA (see section “BGC clustering” above) were annotated using protein domains from Pfam (v35.0; Mistry et al. 2021) as described previously (Larralde et al. 2025). The resulting weighted protein domain compositions were extracted as a sparse matrix for UMAP construction (see section “UMAP construction” below).

### UMAP construction

To visualize the GCF representative set in two-dimensional space, a UMAP (McInnes, Healy and Melville 2020) was constructed in R (v4.3.1), using the sparse matrix of protein domain compositions as input (see section “Protein domain annotation” above), and the following R packages: Signac (v1.12.0; Stuart *et al*. 2021), Seurat (v5.0.2; Hao *et al*. 2024), reticulate (v1.42.0; https://github.com/rstudio/reticulate) and clustree (v0.5.1; Zappia and Oshlack 2018).

Briefly, the sparse matrix of protein domain compositions was read with the scipy.sparse$load_npz function and converted to an RDS object. The RDS object was converted to a chromatin assay (CreateChromatinAssay()) and thereafter to a SeuratObject (CreateSeuratObject(), with “peaks” selected as the assay option). The data was normalized with RunTFIDF(), followed by the selection of top features by FindTopFeatures() with the min.cutoff option set to “q0”. Singular value decomposition (SVD) was used for dimensional reduction. The top 30 SVD dimensions were supplied as input to RunUMAP(), with reduction set to “lsi”. A nearest-neighbour graph was constructed with FindNeighbours(), with “lsi” reduction selected and 30 UMAP dimensions included. To select the clustering resolution, a clustree was generated for resolutions between 0.1-0.9 with 0.05 intervals (Supplementary Figure S5). Thereafter, clustering was performed with FindClusters(), with the options algorithm = 3 and resolution = 0.55. Concatenated metadata, derived from BTyperDB as well as the IGUA output, was attached with AddMetaData() (Supplementary Table S1). The metadata also included biosynthetic classes of the GCF representatives, extracted from the GenBank files. If more than one biosynthetic class was present in a GenBank file, the GCF representative was assigned a “Mixed” label (Supplementary Table S1). The biosynthetic classes of MIBiG BGCs were extracted from the accompanying .json file.

### Maximum-likelihood phylogeny creation

A *B. pseudomycoides* maximum-likelihood phylogeny was created from the genomes labelled in BTyperDB as GTDB species *Bacillus_A pseudomycoides*. Also included was the *B. bingmayongensis* type strain genome (BTyperDB ID BTDB_2022-0000619.2) as an outgroup. Initially, .gff files were created from the .fna files by running Prokka (v1.14.6; Seemann 2014) with the –addgenes option in addition to default settings. Thereafter, .gff files were supplied as input to Panaroo (v1.3.4; Tonkin-Hill et al. 2020), and used to create a core gene alignment with default settings and the following options: the clean mode set to strict (--clean-mode strict), a core threshold of 0.95 (--core_threshold 0.95), invalid genes removed (--remove-invalid-genes), the aligner set to MAFFT (--aligner mafft) and a protein family sequence identity threshold of 0.5 (-f 0.5). Constant sites were filtered and counted using snp-sites (v2.5.1; Page et al. 2016) and the -c and -C options, respectively. The resulting core SNP alignment was subsequently used to construct a maximum likelihood phylogeny using IQ-TREE (v2.2.5; Minh et al. 2020) and the following options: a set seed (--seed 1001), one thousand ultrafast bootstrap replicates (-B 1000; Thi Hoang *et al*. 2017), the number of constant sites from snp-sites supplied as an ascertainment bias correction (-fconst 864972,366217,531353,754415), the nucleotide substitution model set to the general time reversible (GTR) model with invariable sites and free rates (GTR+I+R; Yang 1995; Tavarc 1986; Soubrier *et al*. 2012) and the *B. bingmayongensis* type strain genome supplied as an outgroup (-o BTDB_2022-0000619.2). The initial tree contained three outliers (BTyperDB IDs BTDB_2022-0001360.2, BTDB_2025-0003963.1 and BTDB_2022-0002431.2; Supplementary Figure S6), which were removed prior to repeating the steps with Panaroo, snp-sites and IQ-TREE as described above.

### GCF exploration

BGC alignment figures were generated using LoVis4u (v0.1.5; Egorov and Atkinson 2025), with the options -hl -scc --reorient_loci --run-hmmscan --show-all-feature-labels --use-filename-as-id. For the reduced figure, the option --set-group-colour-for conserved was also used and the --use-filename-as-id option was not included. For LoVis4u figures, only BGCs detected by GECCO and/or from MIBiG were included. Individual genes within BGCs were also used as input for BLASTp using NCBI’s online portal (Sayers *et al*. 2024), with default settings used. For GCF0001397, a custom BLASTp database was generated based on the MIBiG (v3.1; Terlouw *et al*. 2023) records for sevadicin (BGC0000426), mycosubtilin (BGC0001103), enterobactin (BGC0000343, BGC0002476), taxlllaid (BGC0001133) and bacillibactin (BGC0002695), with command line BLAST (v2.14.1; Camacho *et al*. 2009) and the makeblastdb command (makeblastdb -dbtype prot). Using this custom BLASTp database, a GCF0001397 representative BGC was used as a query to search for potential matches to the known BGCs using the blastp command, with the columns in the output specified (-outfmt “6 qseqid sseqid pident length mismatch gapopen qstart qend sstart send evalue bitscore qcovs qcovshsp”) and other settings remaining as defaults.

### Data analysis and visualization

Exploratory analyses (e.g. calculating means, maximum and minimum values) were conducted in R (v4.3.1) with the tidyverse package (v2.0.0; Wickham *et al*. 2019). Statistical tests were run using the kruskal.test() function and the dunnTest() function (with method = “holm”) from the package FSA (v0.10.0; https://github.com/fishR-Core-Team/FSA). Plots were generated with ggplot2 (v3.4.4; Wickham 2016), ggtree (v3.10.0; Xu *et al*. 2022), clustree (v0.5.1; Zappia and Oshlack 2018), VennDiagram (v1.7.3; Chen and Boutros 2011), ggridges (v0.5.6; https://github.com/wilkelab/ggridges), ggpubr (v0.6.0; https://github.com/kassambara/ggpubr), ggplotify (v0.1.2; https://github.com/GuangchuangYu/ggplotify), ape (v5.7-1; Paradis and Schliep 2019), and phytools (v2.3-0; Revell 2024).

### Web application construction

The BTyperBGC web application was built in R (v4.3.1) with the Shiny package (v1.8.1.1; https://github.com/rstudio/shiny). The following R packages are used when running the application: tidyverse (v2.0.0; Wickham *et al*. 2019), ggplot2 (v3.4.4; Wickham 2016), DT (v0.33; https://github.com/rstudio/DT) and markdown (v1.12; https://github.com/rstudio/markdown). The application is hosted at https://github.com/c20josbl/btyperbgc_shiny.

### Data availability

BTyperDB and NCBI accession numbers for all genomes used in this study are provided in Supplementary Table S1. GCF clustering results are available in Supplementary Table S1.

### Code availability

BTyperBGC is open-source (GPL-3.0 license) and available at https://github.com/c20josbl/btyperbgc_shiny.Computational analyses are described in the manuscript; code is additionally available at GitHub: https://github.com/c20josbl/btyperbgc_manuscript_documentation.

## ACKNOWLEDGMENTS

JB, HG, VR, and LMC were supported by the SciLifeLab & Wallenberg Data Driven Life Science (DDLS) Program (grant: KAW 2020.0239). Additional funding was provided by the Swedish Research Council (VR grants 2023-05212, 2024-03952, 2024-06085), and the Swedish Cancer Society (grant number #233102 Pj). This research was conducted using the resources of High Performance Computing Center North (HPC2N). Additional computations/data handling were enabled by resources provided by the National Academic Infrastructure for Supercomputing in Sweden (NAISS), partially funded by the Swedish Research Council through grant agreement no. 2022-06725.

## AUTHOR CONTRIBUTIONS

Computational analyses were performed by JB, with input from HG, ML, JH, and LMC. Software development was performed by JB, with input from HG, JH, and LMC. Genomic data and associated metadata were provided/curated by VR and LMC. LMC conceived and funded the study. JB and LMC co-wrote the manuscript with input from all authors.

## SUPPLEMENTARY FIGURES

**Supplementary Figure S1.**
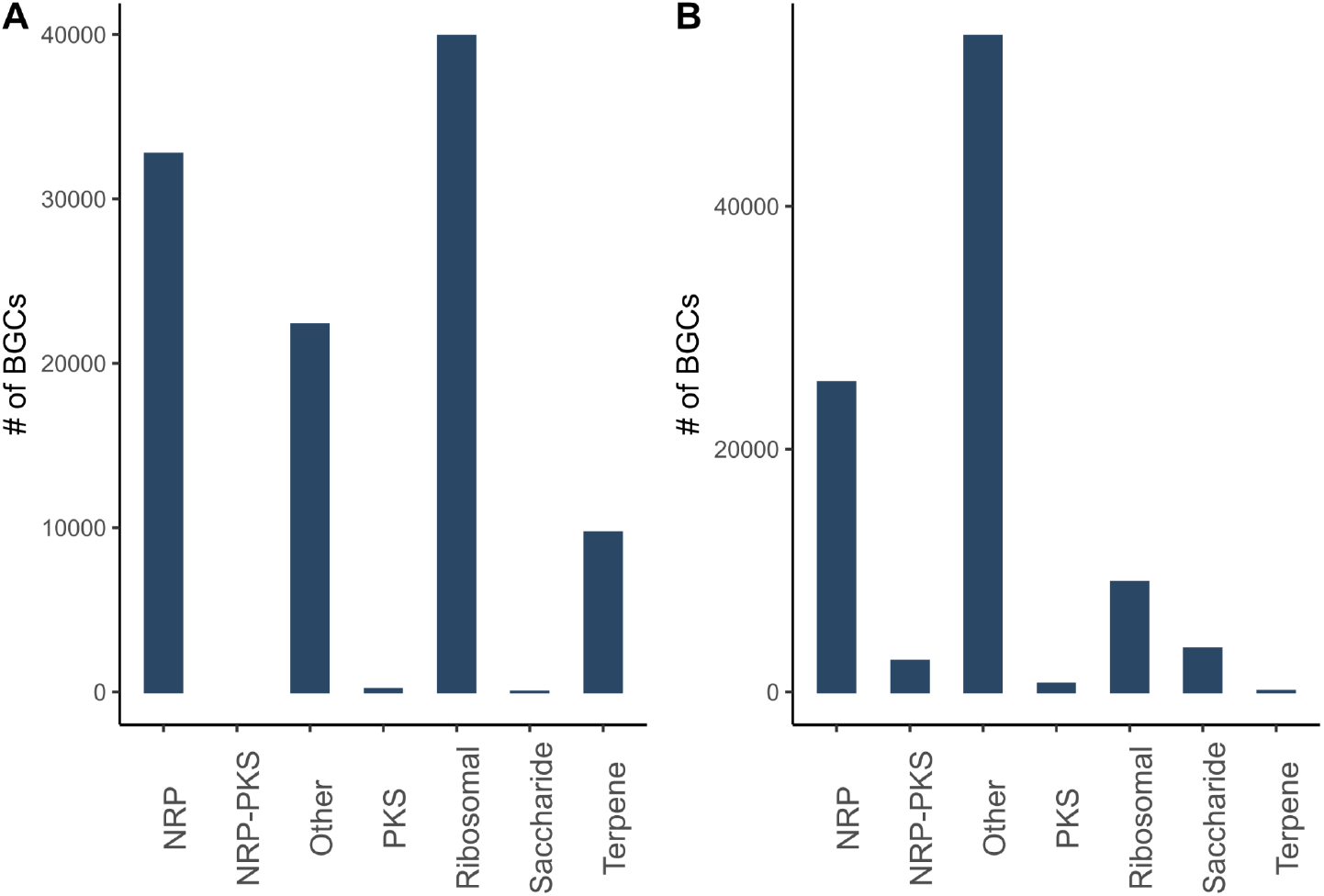
Chemical class distribution of BGCs detected by (A) antiSMASH and (B) GECCO.

**Supplementary Figure S2.**
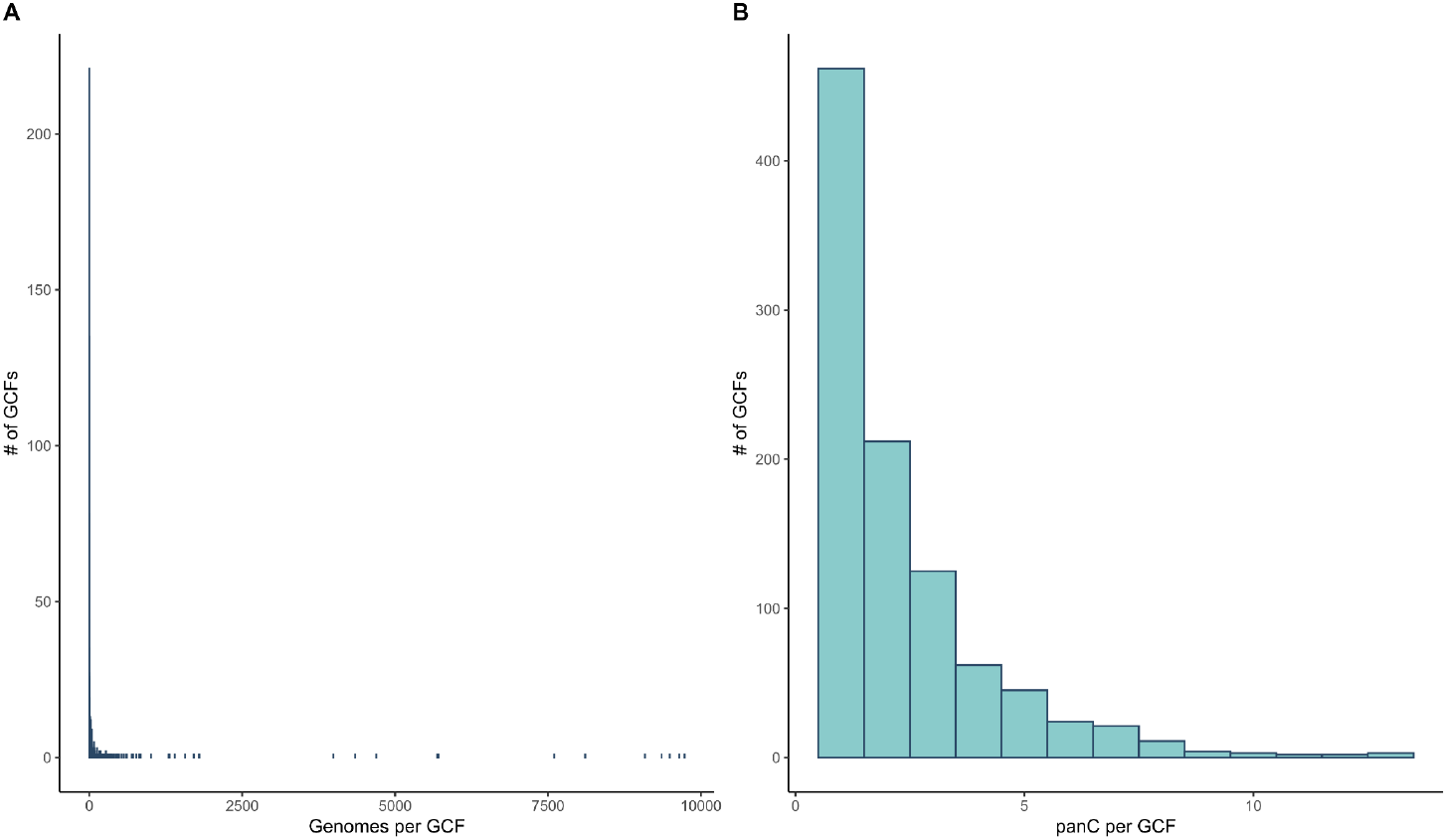
(A) Histogram of the number of genomes (x-axis) per GCF (y-axis) (Bl Histogram of the number of *panC* groups (x-axis) per GCF (y-axis). For both (A) and (B), no MIBiG BGCs were included.

**Supplementary Figure S3.**
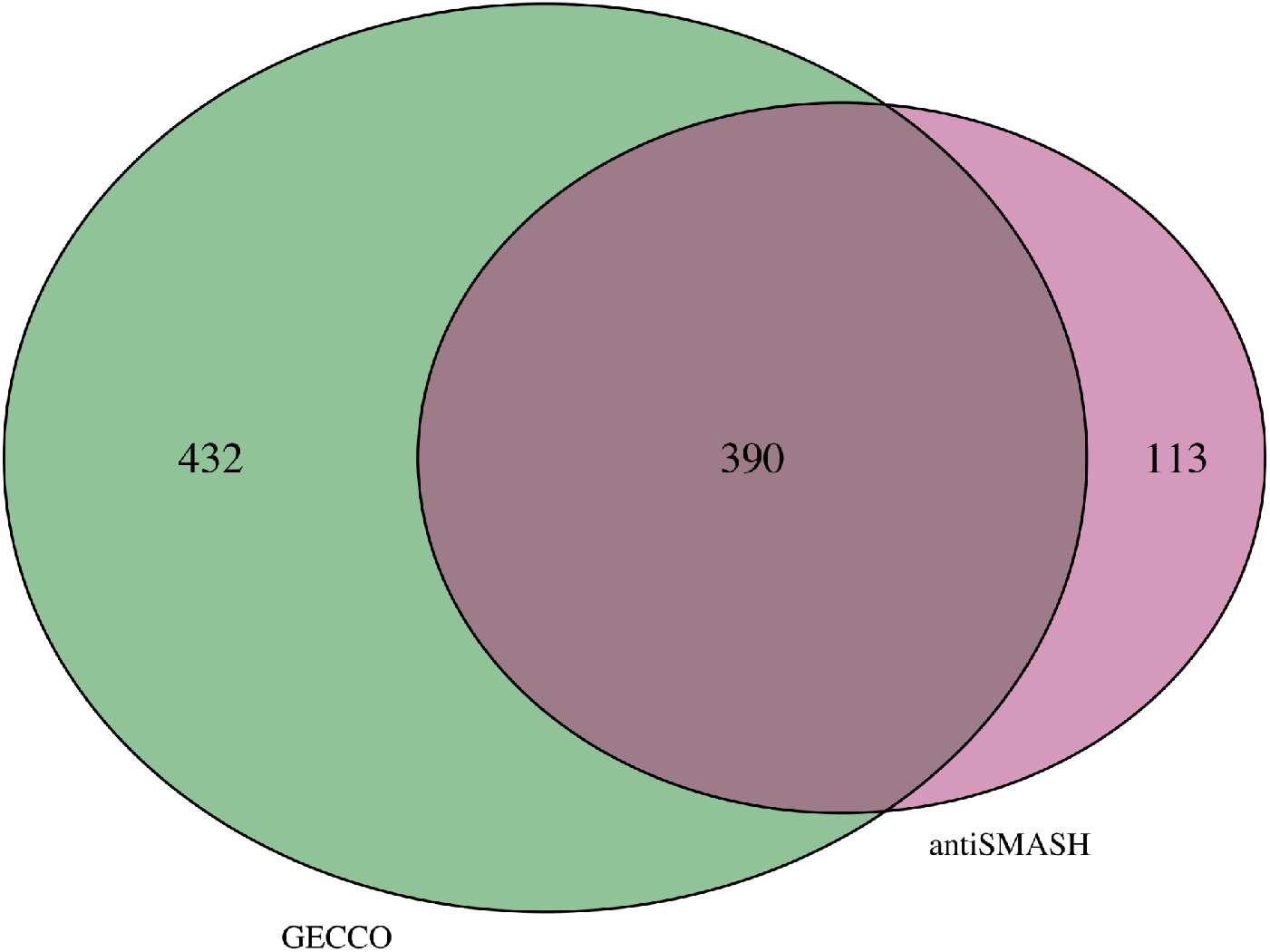
A Venn diagram showing the composition of gene cluster families (GCFs) absent of MIBiG BGCs. The overlap represents GCFs which include BGCs derived using both GECCO and antiSMASH.

**Supplementary Figure S4.**
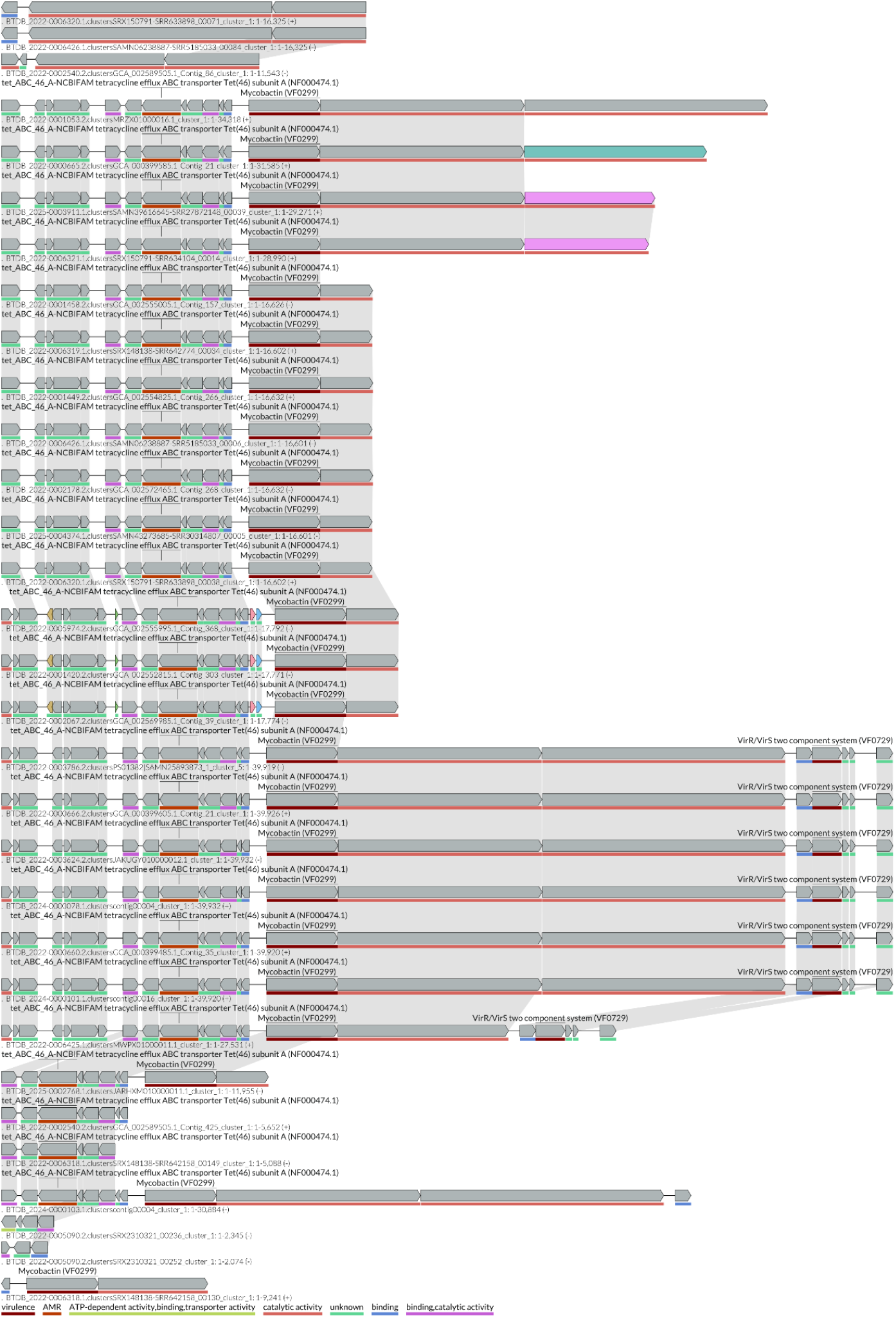
LoVis4u plot of GCF0001397.

**Supplementary Figure S5.**
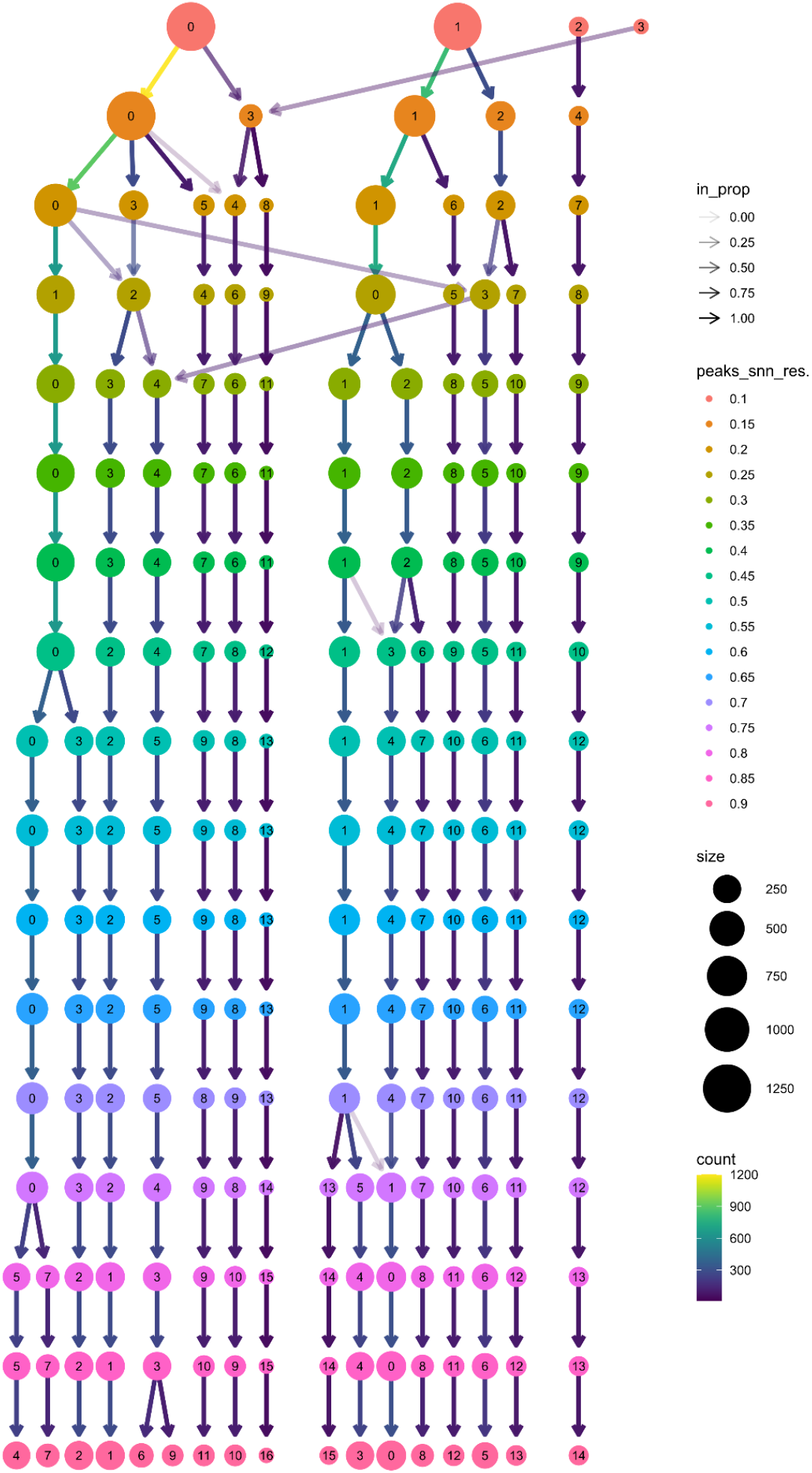
Clustering tree for the BTyperBGC UMAP.

**Supplementary Figure S6.**
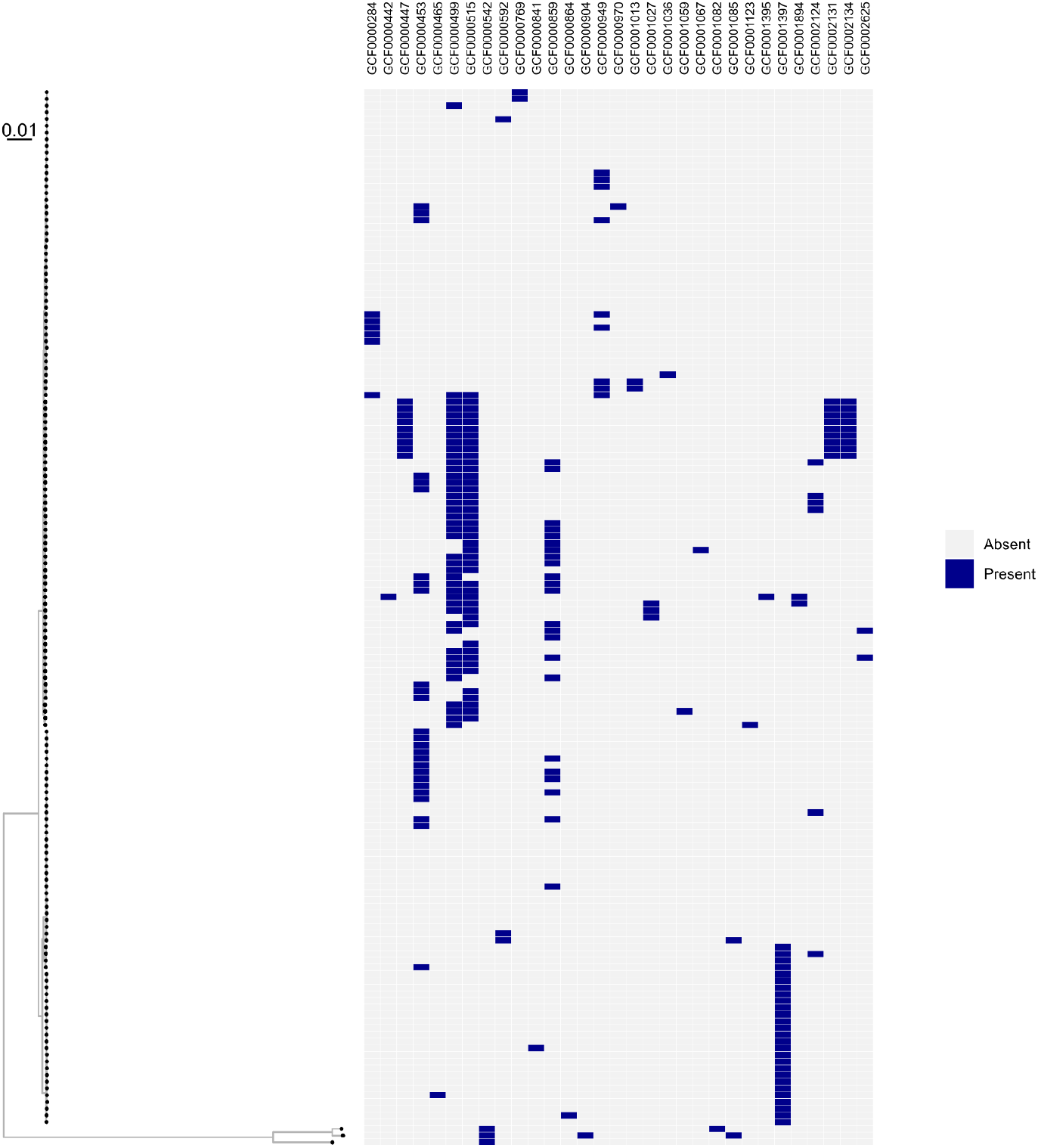
Maximum-likelihood phylogeny of all GTDB species *Bacillus pseudomycoides* genomes (left), annotated based on presence/absence of BGCs in *B. pseudomycoides*-unique que GCFs (right). Branch lengths represent substitutions per site. The phylogeny was rooted via an outgroup (*B. bingmayongensis* type strain genome; BTyperDB ID BTDB_2022-0000619.2).

